# biPACT: a method for three-dimensional visualization of mouse spinal cord circuits of long segments with high resolution

**DOI:** 10.1101/2021.12.12.472305

**Authors:** Katsuyuki Nakanishi, Munehisa Shinozaki, Narihito Nagoshi, Masaya Nakamura, Hideyuki Okano

## Abstract

**Background:** The spatial complexity of neuronal circuits in the central nervous system is a hurdle in understanding and treating brain and spinal cord injuries. Although several methods have recently been developed to render the spinal cord transparent and label specific neural circuits, three-dimensional visualization of long segments of spinal cord with high resolution remains challenging.

**New Method:** We present a method that combines tissue staining of neuronal tracts traced with biotinylated dextran amine (BDA) and a modified passive clarity clearing protocol to describe individual fibers in long segments of mouse spinal cord.

**Results:** Corticospinal tract was traced with BDA with a mouse model of thoracic spinal cord injury. The spinal cord was stained and cleared in two weeks with four solutions: staining solution, hydrogel solution, clearing solution, and observation solution. The samples were observed with a light-sheet microscope, and three-dimensional reconstruction was performed with ImageJ software. High resolution-images comparable with tissue sections were obtained continuously and circumferentially. By tiling, it was possible to obtain high-resolution images of long segments of the spinal cord. The tissue could be easily re-stained in case of fading,

**Comparison with Existing Methods:** The present method does not require special equipment, can label specific circuits without genetic technology, and re-staining rounds can be easily implemented. It enables to visualize individual neural fiber of specific neural circuit in long spinal cord segments.

**Conclusions:** By using simple neural staining, and clearing methods, it was possible to acquire a wide range of high-resolution three-dimensional images of the spinal cord.

**Highlights:**

- No special devices or genetic tracers are required for a new clearing method
- Neuronal fibers are individually depicted in long segments of mouse spinal cord.
- Re-staining of neuronal fiber is possible.
- Stereotaxic observation is achieved by 3-D reconstruction with open-source software.

## 1. Introduction

Every year, approximately 0.93 million people suffer from spinal cord injury (SCI)(Lee et al., 2014). Since the ability of central nervous system (CNS) neurons to recover in adult mammals, including humans, is extremely limited, many patients have their daily lives severely affected due to sequelae, resulting in massive social losses. In recent years, various regenerative approaches have seemed to induce some level of neural network reconstruction at the injured site(Hilton and Bradke, 2017; Ramer et al., 2014). To further advance the regenerative treatment of neural circuits, it is crucial to accurately assess the condition of the remaining neuronal tracts after SCI and their potential change upon treatment. Although tissue evaluation using slice sections is the method commonly used to evaluate neural circuits, spinal cord neural circuits form complex three-dimensional networks(Kim et al., 2013; Steward et al., 2003; Tian and Li, 2020; Vigouroux et al., 2017). Therefore, it is essential to perform three-dimensional analyses at a resolution that can trace individual fiber to accurately assess the extent of damage and the possible effect of a given treatment(Erturk et al., 2011; Hilton et al., 2019).

The recent development of light-sheet microscopy, together with various methods for clearing tissues, has made it possible to observe tissues without sectioning(Chung et al., 2013; Murakami et al., 2018; Tainaka et al., 2016; Treweek et al., 2015; Ueda et al., 2020; Yang et al., 2014; Zhu et al., 2019). By making the neural tissue transparent after fluorescent labeling of a specific circuit or tissue and observing it under a microscope equipped with objective lenses of long working distance, it is possible to visualize the three-dimensional structure in situ. However, to introduce transparency methods, it is necessary to solve the following problems. 1) Special device and/or long time necessary to render the spinal cord transparent. 2) Mismatch between transparency and the observation method. 3) The requirement of genetically modified mice or viruses for fluorescent labeling. 4) Reduction of fluorescence (fading) during the staining and clearing procedure. 5) Incompatibility between the resolution to follow a single neural fiber and the imaging of wide area of CNS.

To solve these problems, we used traditional and inexpensive biotinylated dextran amine (BDA) modifying passive clarity technique (PACT)(Bouvier et al., 2016; Neckel et al., 2016; Woo et al., 2016; Yu et al., 2017), and established a method biPACT, that requires neither special devices such as vacuum devices nor genetic manipulation. Clearing was completed while maintaining fluorescence simply by exchanging the solution, allowing three-dimensional observation to be performed with the same resolution as sectional tissue, which enables to trace individual neuronal fibers.

## 2. Materials and Methods

The flow of the experiments is shown in Figure 1. A SCI model mouse is created and a classical monosynaptic neuronal tract tracer, BDA, is injected into the sensorimotor cortex of the cerebrum. Once the tissue is collected, it is observed with a light-sheet microscope after a 2-week staining and clearing process.

**Figure 1.**
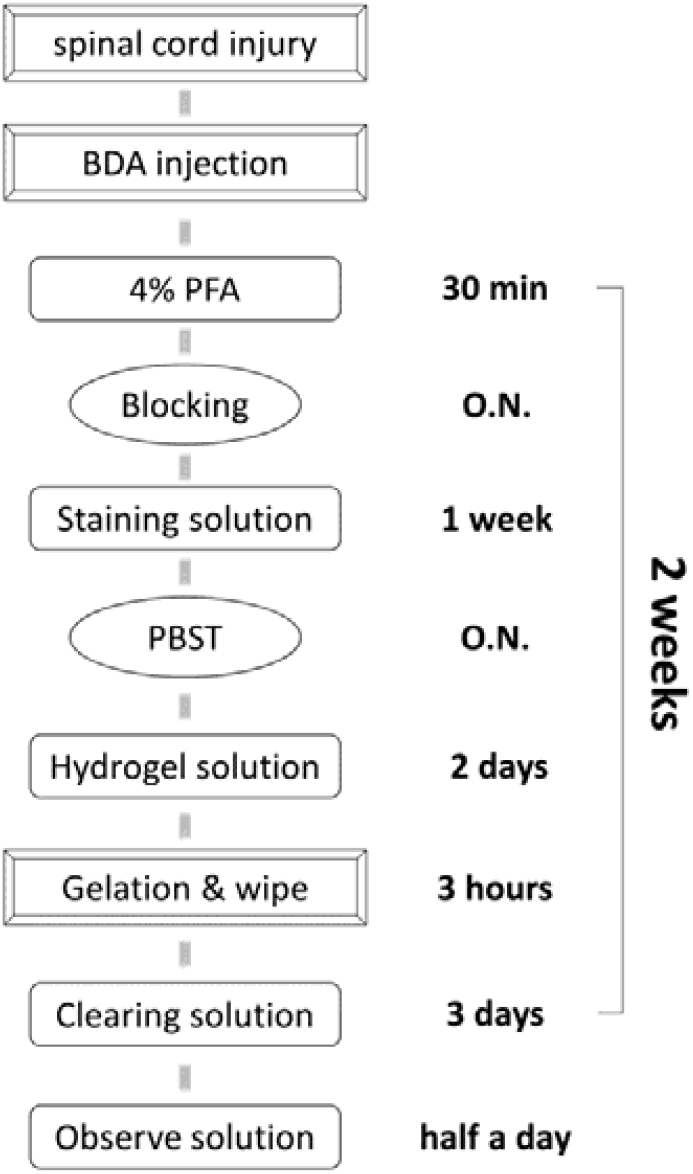
Process flowchart. The flow from spinal cord injury to observation is shown. The entire process from sample collection to imaging takes two weeks. Induction of spinal cord injury is followed by injection of BDA into the sensorimotor cortex. After removing the spinal cord, post-fixation is performed with PFA at room temperature for 30 minutes, and the spinal cord is immersed in the blocking solution overnight at 4°C. Then the tissue is stained in the staining solution at 4°C for one week. After washing overnight with PBST at 4°C, the tissue is transferred to hydrogel solution and immersed at 4°C for two days. The tissue is incubated in a water bath at 37°C for 3 hours, and the gel around the tissue is gently wiped. After that, it is transferred to a thermostat chamber at 47°C to make it translucent. When the tissue becomes translucent, it is transferred it to an observation solution at room temperature and it becomes completely transparent after half a day.

### 2.1. SCI model

All experiments were carried out according to the Guidelines for the Care and Use of Laboratory Animals of Keio University (Assurance No. 13020) and the Guide for the Care and Use of Laboratory Animals (NIH). Eight-week-old black6/J female mice were anesthetized with an intraperitoneal injection of ketamine (100 mg/kg) and xylazine (10 mg/kg), and a contusive SCI was induced at Th10 using an IH impactor (70 kdyn; Precision Systems and Instrumentation) as previously described(Shibata et al., 2021).

### 2.2. Anterograde labeling of the corticospinal tract (CST)

Six weeks after injury, biotinylated dextran amine (D1956, BDA; MW 10,000; 10% in distilled water, Invitrogen, USA) was injected into the sensorimotor cortex at four sites (coordinates: 0.5–1.5mm posterior to bregma and 0.5–1.5mm lateral to bregma) to trace descending CST fibers for hindlimbs, as previously described(Cafferty et al., 2010). Two weeks later, the mice were deeply anesthetized and transcardially perfused with 4% paraformaldehyde (PFA), and the spinal cords were removed.

### 2.3. Devices and Solutions

The devices and solutions used are shown in Figure 2A. One solution is needed for staining, and three solutions, a hydrogel solution, a clearing solution, and an observation solution, are required for clearing. Reagents for the preparation of the hydrogel solution should be cooled in advance, mixed on ice, and the resulting solution should be dispensed into 50 ml conical tubes that can be stored at -20°C, and thawed at 4°C before use. The clearing solution can be prepared and stored at room temperature; NaOH is added to bring the pH to 8.5. The observation solution can be prepared and stored at room temperature. Sorbitol is difficult to dissolve, but it dissolves naturally when left at room temperature.

**Figure 2.**
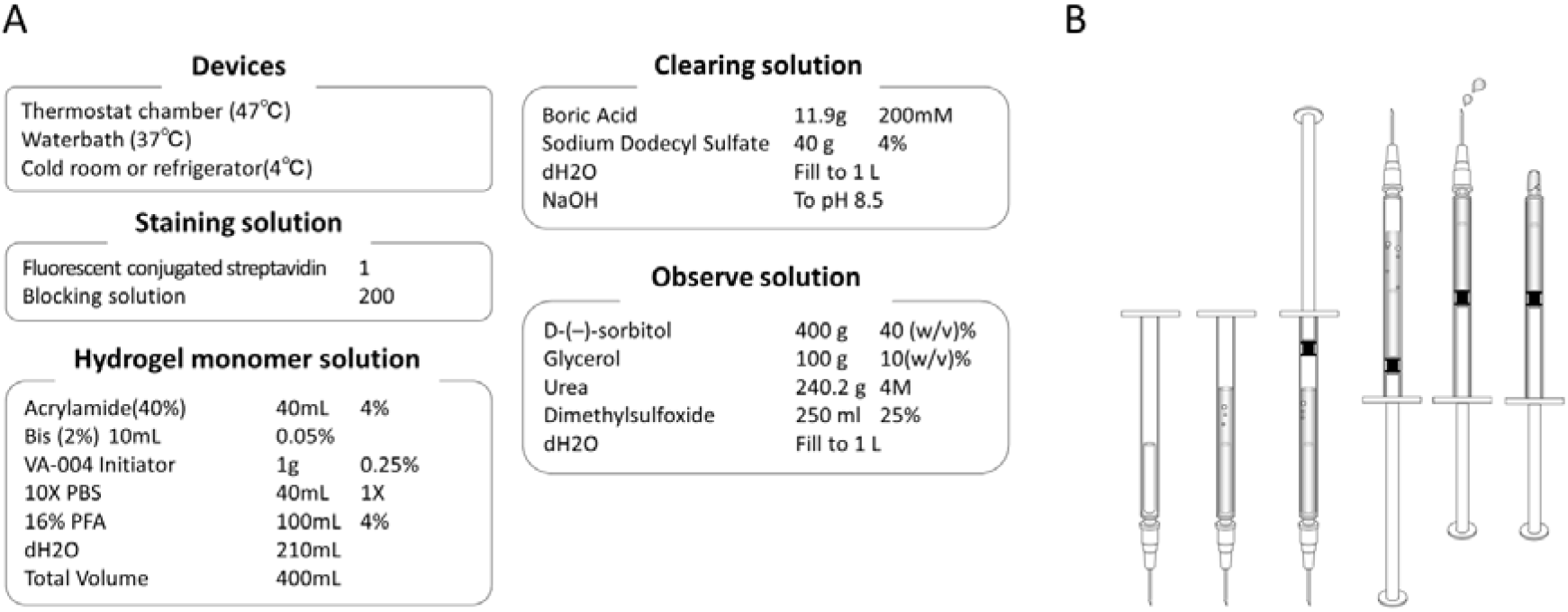
Tools required for transparency and gelation procedure. The devices and solutions required for the biPACT are shown in A. The required devices are a refrigerator, a 37°C water-bath, and a thermostat chamber that can be set to 47°C. Fluorescence-conjugated streptavidin is diluted 1: 200 with a blocking solution. Hydrogel solution is mixed on ice and stored frozen until use. It is thawed at 4°C the day before use. The clearing solution is mixed at room temperature. The observation solution is also mixed at room temperature. The procedure for gelation step is shown in B. No vacuum equipment or nitrogen gas is required. Attach a thin needle to a 1 ml syringe and remove the plunger. Put the tissue inside and fill with hydrogel solution. The plunger is returned, and the syringe is turned the tip up. While tapping, the plunger is pushed little by little to remove the air bubbles inside. When hydrogel solution comes out from the needle tip, the needle is removed, and the tip is sealed with parafilm to prevent air from entering.

#### 2.3.1. Staining step

Once dissected from the spine, the spinal cord (SC) was soaked in 4% PFA at room temperature for 30 minutes. The dura mater was then peeled using two sharp-pointed tweezers and the SC was immersed overnight in blocking solution at 4°C. Staining solution is constituted of Alexa Fluor 555 Streptavidin (S21381, Thermo Fisher Scientific, USA) and usual blocking solution (03953-95, Blocking One, Nacalai, Japan). The SC was then soaked in staining solution (1 ml per cm of spinal cord tissue) at 4°C for seven days, under optional orbital shaking. Intense staining was obtained with a dilution of 1: 200, but if the background is strong, it is recommended to work with lower dilutions. The SC was protected from light during the overnight washing step in PBS-T (5 ml for 1 cm spinal cord tissue) at 4°C. At the same time, the thawing of the hydrogel solution at 4°C was initiated. The next day, the SC was transferred to the hydrogel solution (5 ml for 1 cm of spinal cord tissue) for two days at 4°C, protected from light, with possible orbital shaking.

#### 2.3.2. Gelation step

The plunger of a 1 ml syringe was removed, and a thin (≥ 26G) metal needle was attached to the syringe (Figure 2B). The SC was placed in the lumen of the syringe, and the hydrogel solution was added. Once the plunger was attached, the syringe was flipped with its tip facing upward. Gentle tapping of the syringe removed air bubbles. The plunger was then pushed until all air came out of the tip of the needle. The needle was removed and parafilm was used to hermetically seal the syringe. Then the syringe protected from light with aluminum foil was incubated in a 37°C water bath for 3 hours. A thermostat chamber is not adaptable instead of the water bath. At the end of the gelation step, the plunger was removed and the internal gel and the SC were collected on a clean surface of paper towel. The gel around the SC was gently but thoroughly removed with a thin paper towel.

#### 2.3.3. Clearing step

The SC was then transferred to a tube containing the clearing solution (10 ml for 1 cm of spinal cord tissue). Protected from light, this tube was placed in a 47°C thermostatic chamber. The SC was examined every day, which allowed us to notice that the mouse spinal cord became translucent in three days. The clearing period can be extended as an option. It was possible to delay the observation for up to two weeks if the sample is protected from light and stored at 4°C. Alternatively, when using a lens with an RI of 1.38, it is possible to skip the next RI matching step and use the clearing solution to use as an observation solution. In that case, it takes ten days before the SC becomes transparent in the clearing solution from translucent condition.

#### 2.3.4. Refractive Index (RI) matching step

Half a day before microscopic observation, the SC were transferred to an observation solution (5 ml for 1 cm of spinal cord tissue), protected from light, and kept at room temperature. This made the SC completely transparent. Since fading progresses quickly in the observation solution, observing within three days after transfer to the observation solution is preferable. If the fluorescent staining of the cleared tissue has faded, re-staining can be performed by soaking the SC in blocking solution overnight, followed by the staining and washing procedure.

### 2.4. Observation and reconstruction

A light-sheet microscope (Zeiss Z1 microscope; Carl Zeiss, Göttingen, Germany), equipped with a 5× lens with an RI of 1.45, was used. The observation chamber was filled with the observation solution and a holder was used to maintain the transparent spinal cord in the center of the chamber. A continuous image was captured with the optimal interval condition in single-side mode. Three-dimensional images were obtained by reconstructing images using the 3D-project plugin of ImageJ software (NIH. 64-bit Java 1.8.0, United states).

### 2.5. Two-dimensional histological evaluation

Spinal cord frozen sections were prepared using a Leica cryostat (Leica CM3050 S, Leica Microsystems, Wetzlar, Germany) to confirm the fluorescence staining. For preparing frozen sections of the clarified tissue, the SC was first immersed in 4% PFA, followed by overnight incubation in a 20% sucrose solution. A series of 20-μm spinal cord axial sections was prepared and imaged with a fluorescence microscope (BZ-9000, Keyence, Japan).

## 3. Results

### 3.1. Clearing

Sections prepared with the stained SC confirmed that the BDA-traced CST was intensely labeled over time (Supplementary Figure 1). When the spinal cord in the hydrogel solution was incubated at 37°C for 3 hours, the solution around the spinal cord also got gelled, suggesting that the spinal cord was sufficiently polymerized (Fig. 3A-C). In the clearing step, the margins of the spinal cord tissue were gradually cleared over time in 47°C chamber (Fig. 3D), and a translucent tissue was obtained on the third day (Fig. 3E). The refractive index matching solution had the SC tissue completely transparent (Fig. 3F).

**Figure 3.**
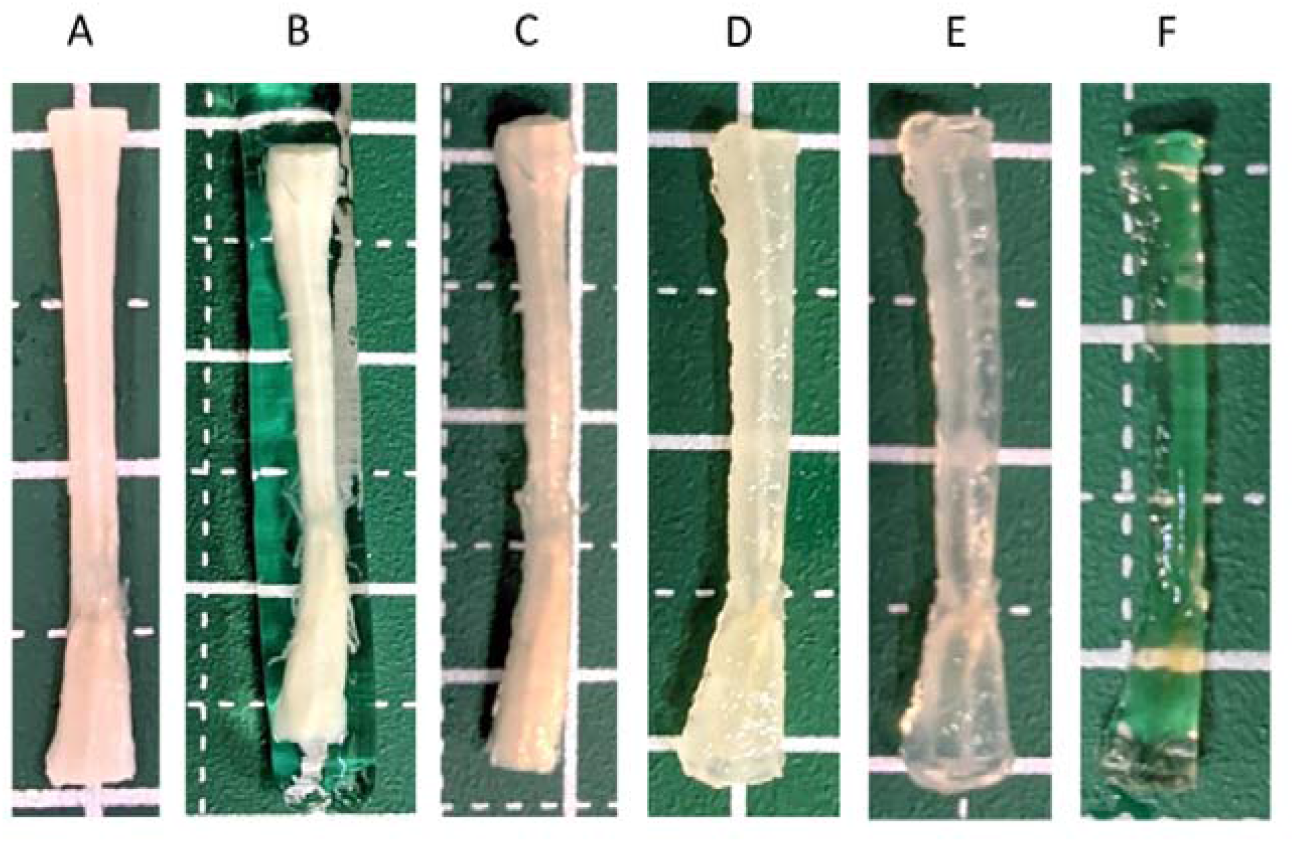
Tissue transparency process. The process of tissue transparency is shown in Figure3. (A) Tissue after soaking in hydrogel solution. (B) The tissue after the gelation was wrapped in gel when removed from the syringe, suggesting that the hydrogel solution that permeated the inside was sufficiently polymerized. (C) Tissue from which the gel has been gently removed. (D) The tissue on the second day during the clearing step. (E) The tissue on the third day during the clearing step. It is still translucent on the third day, and it is sufficient to move on to the next step. (F) The tissue soaked in observation solution for half a day is completely transparent.

### 3.2. Observation

Imaging of the labeled CST was performed with a light-sheet microscope. The observation was carried out in three directions chosen to be orthogonal to the anatomical axes of the spinal cord, and clear images were obtained at all shooting angles (Figs. 4A-C and Supplementary Movies 1-3). In the axial direction, in addition to white matter fibers, fibers in grey matter and their varicosities could be visualized, demonstrating that sufficient resolution was obtained. When vertically tiling, light was emitted parallel to the long (rostro-caudal) axis of the spinal cord, allowing observation of the spinal cord over a long range of 868 μm (Fig. 4D and Supplementary Movie 4) with the same resolution.

**Figure 4.**
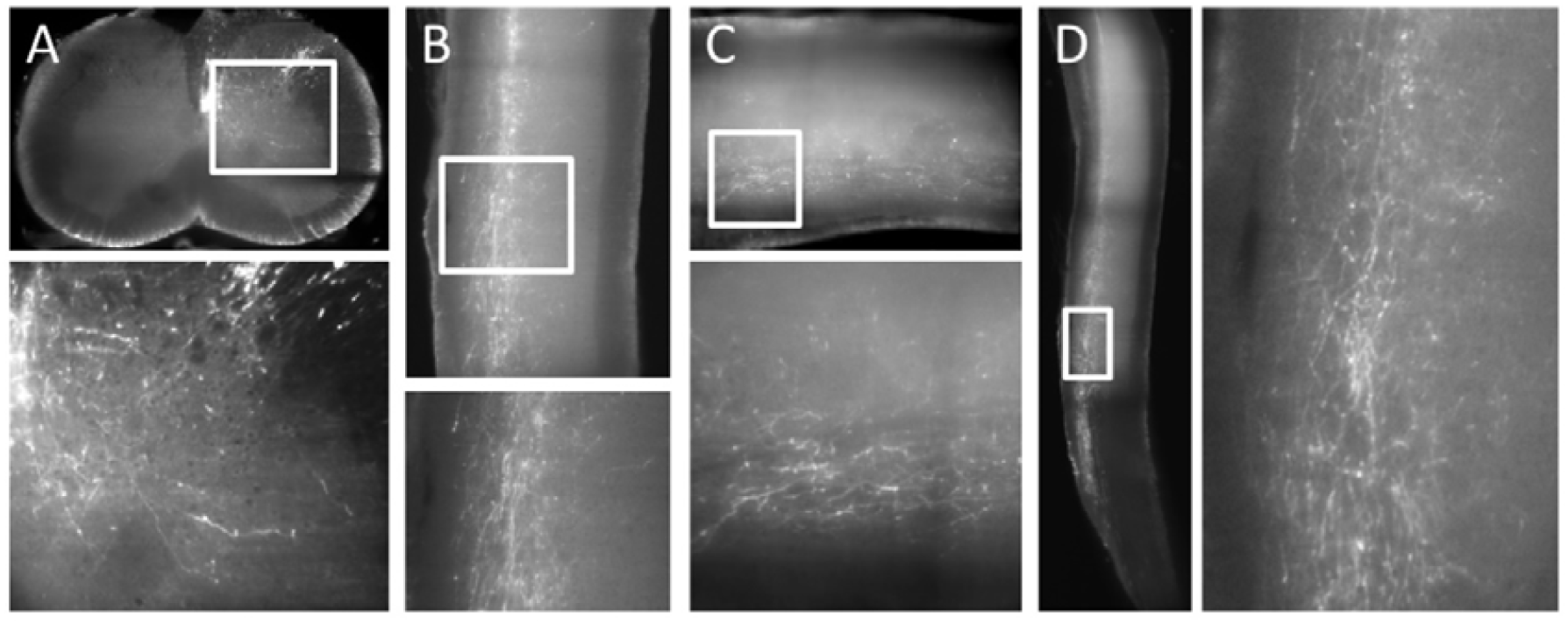
Images acquired with a light-sheet microscope. A representative image of tissue taken continuously from three directions with respect to the axis of the spinal cord. The imaging conditions are as follows: magnification 0.5, 1920 × 1920 pixels, pitch of 2.27. See Supplementary Movies 1-3 for a series of images of the entire spinal cord displayed here. In the axial (A), sagittal (B) and coronal (C) axes, the resolution obtained allowed one to visualize individual fibers. In the axial axis, the varicosity of the fibers in the gray matter can be visualized. (D) Tiled image in the sagittal direction to obtain a long-segment view of the spinal cord. Tiling in the sagittal direction can be done with constant brightness because the optical path distance in the tissue does not change. See Supplementary Movie 4 for the entire series of consecutive images.

### 3.3. Three-dimensional reconstruction

A three-dimensional image was reconstructed from the sequential images using ImageJ software (Fig. 5A and Supplementary Movie 5). When taken at higher magnification followed by horizontal tiling, it was possible to depict the fine fibers that enter the gray matter from the white matter of the spinal cord (Fig. 5B and Supplementary Movie 6). Additionally, the vertically-tiled images showed that the density of the CST varied depending on the segments. It also showed a dislocation of the CST fibers on the cranial side of the lesion, suggesting a sprouting of the residual fibers (Fig. 5C and Supplementary Movie 7).

**Figure 5.**
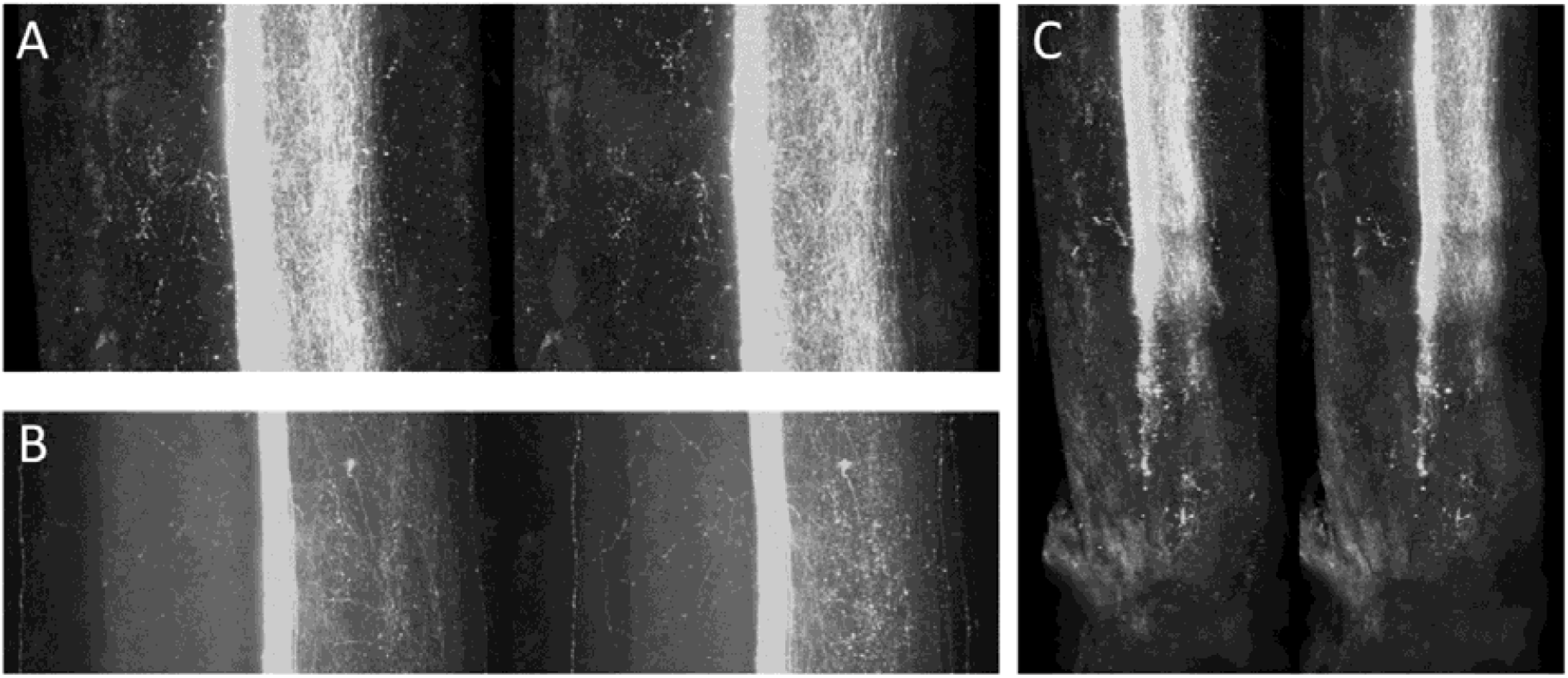
Three-dimensional reconstruction from continuous images. Three-dimensional reconstruction from series of continuous images using the ImageJ 3D-project plugin. Since two images with different angles are arranged side by side, stereoscopic viewing is possible by the crossover method. See Supplementary Movies 5-7 for the rotational movies. (A) Sagittal image. (B) Two sagittal images were captured with high magnification and horizontally tiled to include the entire circumference. An even higher-resolution image was obtained. (C) Vertical tiling image of the spinal cord. A long range of evaluations can be performed at once by tiling in the long axis direction. Note that the density of the corticospinal tract is not consistent in the normal part of the spinal cord. It was demonstrated that the fibers in the corticospinal tract in the white matter decreased, and the fibers in the gray matter were dislocated around the lesioned area.

## 4. Discussion

We used a mouse SCI model to generate a three-dimensional depiction of circuit-specific neural fibers using a classical tracer BDA, a modified PACT, and a light-sheet microscope. Our method does not require viruses, genetically modified animals, or any special clearing device. The final resolution was high enough to visualize precisely up to single fibers. BDA is the most widely used neuronal tracer(Cafferty et al., 2010; Liang et al., 2015; Steward et al., 2008a; Zareen et al., 2018), and it would be meaningful to be able to easily observe BDA tracing in 3D with the same accuracy as 2D-based analyses.

As confocal, two-photon, and light-sheet microscopes were shown to be suitable for 3D analysis of the CNS, several clearing methods have been reported. Still, many factors are limiting the generalized use of such techniques(Richardson and Lichtman, 2015).

### 4.1. Devices and procedures for observation

It is important to select the transparency method depending on the microscope used. When observing a block tissue, the optical path passing through a transparent tissue becomes long, and light is scattered. Therefore, if the laser power is insufficient compared to the level of light scattering, a fine image cannot be obtained. In mild or medium transparency methods such as ScaleS or SeeDB, clear images can be obtained by using a two-photon microscope, whose laser power can be changed according to the depth(Hama et al., 2015; Helmchen and Denk, 2005; Ke et al., 2013; Schiessl and Castrop, 2016; Susaki et al., 2014). Since the laser power of the light-sheet microscope cannot be changed according to the depth, it is necessary to use strong transparency methods such as DISCO system(Erturk et al., 2012; Quinta et al., 2015) or CLARITY(Epp et al., 2015; Jensen and Berg, 2017; Tomer et al., 2014) in which light absorption and scattering are reduced. In the present study, by combining PACT with a high-RI observation solution, we obtained transparency sufficient for observation with a light-sheet microscope in a short period.

The choice of clearing method also includes hardware issues. Zeiss’s light-sheet microscope uses a sample chamber in which the glass is fixed with a rubber that is not compatible with organic solvents. Samples cleared with methods using organic solvents such as DISCO necessitate the use of specific chambers or an alternative observation solution. The present biPACT method is a water-based protocol, and it is much easier to be implemented. In addition, to match the high endogenous RI components is crucial, and an observation solution of high RI is required. In the Zeiss light-sheet microscope, there are three types of RI settings for the objective lens: 1.33, 1.38, and 1.45. In this protocol, we obtained a highly transparent optical path by matching the observation solution and the microscope setting with the RI of 1.45.

### 4.2. Tissue staining and prevention of fading

One of the most important factors for clear observation of neural tissue is the intensity of fluorescence of the labeled neurons. For fluorescent labeling, the most common methods include using transgenic animals that express fluorescence in a specific tissue, injecting viruses to introduce a reporter gene into a target neural circuit, and staining neuronal tracers(Soderblom et al., 2015). Here, we use BDA and fluorescence-conjugated streptavidin to enable clear neural fiber-imaging. BDA does not require genetic manipulation, is stable in tissues for a long period, and delineates fibers in the finest detail. Unlike viruses, its use is not constrained by biosafety concerns and can be injected with a physical pump and a metal needle, which have the advantage of being easy to be introduced into research facilities(Han et al., 2012; Hellenbrand et al., 2013; Steward et al., 2008b).

To perform block staining, it was possible to use the same staining solution used for sections simply by extending the staining duration in the present study. Many methods of immunostaining tissues have been reported along with the clearing method(Belle et al., 2014; Du et al., 2019; Kubota et al., 2017; Lee et al., 2016; Martinez-Lorenzana et al., 2021; Renier et al., 2014), and some even label RNA within the clearing tissues(Sylwestrak et al., 2016; Wang et al., 2018a; Ye et al., 2016). However, the appropriate conditions must be precisely determined for almost every antibody. During this study, we put a lot of effort into optimizing immunostaining, but we were unable to obtain satisfactory results (data not shown), even when we significantly extended the incubation period of antibodies. The technical refinements would solve this issue in the future.

Fluorescence-conjugated streptavidin could intensely stain the deepest part of the mouse spinal cord in one week. Several key differences from usual immunostaining may explain such contrasting results. The protocol is one-step staining, and streptavidin has a low molecular weight compared to general secondary antibodies. The 60 kD-fluorescence-conjugated streptavidin can penetrate the SC relatively easily, and the unbound streptavidin can be washed equally easily after staining. The high intensity of the signal might be achieved by the binding of multiple streptavidin molecules to biotin. The stronger binding between streptavidin and biotin (Kd = 10^−14^ mol/L), compared to the classical affinity of antigen-antibody (Kd = 10^−6^ to 10^−9^ mol/L), also could contribute, at least theoretically, to the enhanced signal/noise ratio.

The second factor necessary for observing the fiber tracts within the cleared tissue is the reduction of fading of fluorescence that results from the clearing process. Generally, fading is usually more pronounced when the clearing stringency increases. For example, the original Clarity method strongly eliminates lipids(Chung et al., 2013). The DISCO-based method, which uses an organic solvent, easily fades fluorescent proteins such as the Green Fluorescent Protein (GFP)(Tian and Li, 2020). For those methods, finely tuning of conditions to balance the final degree of transparency with the remaining fluorescence is required. Our protocol has relatively wide range of the optimal conditions for clearing and retaining fluorescence.

The evaluation of the brain circuits is also needed for the research of SCI. The distance from the tissue surface to the most internal areas is longer in the brain, implying that the fluorescence present at the surface may fade before the deep part becomes clear. Improvements of the protocol would be needed for the brain analysis.

### 4.3. High-resolution observation

As the name implies, light-sheet microscopy is based on generating a sheet of light within the specimen and ensuring that it coincides with the focal plane of a high numerical aperture objective placed perpendicularly(Hillman et al., 2018; Migliori et al., 2018). Therefore, even in the 1920 × 1920 range, only one exposure time is required. If this is a confocal microscope, an image is formed while scanning each point, so it takes several tens of times longer observation time. In this study, it took about 5 minutes to capture the mouse spinal cord around the circumference at a resolution of 1 µm / pixel, and very fast imaging was possible. However, if the tissues are not transparent enough, the excitation light scatters, and the image inside the tissue becomes very blurry(Ueda et al., 2020). Even if the transparency was obtained such that characters printed on a paper could be discerned, it was not possible to observe it with a light-sheet microscope (data not shown). Our empirical observations suggest that accurate imaging with a light-sheet microscope requires a level of transparency sufficient to lose sight of the tissue immersed in the observation solution.

In addition, the light-sheet microscope captures images while translating the excitation plane in the Z-axis direction to obtain continuous images. The excited plane itself is thick, which reduces the risk of overlooking fluorescence and is suitable for tracking of individual fibers. On the other hand, the thickness is coarser than the pitch of the XY plane, and the thickness of the excitation plane differs between the center and the edge of the field of view(Remacha et al., 2020), so the resolution in the Z-axis direction further decreases. Therefore, unlike confocal microscopes and two-photon microscopes, it is necessary to carefully reconstruct the images obtained with the light-sheet microscope into three-dimensional coordinates before quantifying them. For example, when capturing a long area of the spinal cord, it is advantageous to shoot the excitation light parallel to the long axis. But in this case, it is difficult to accurately quantify fibers in the reconstructed axial plane. In the future generation of light-sheet microscopes, it is expected that the thickness of the excitation light becomes thinner and remains constant between the center and the edge of the field. In addition, it is also desirable to develop software that removes image data, which overlap in the Z-axis direction and interfere with concatenating operations.

We used ImageJ, NIH’s free software, for the three-dimensional reconstruction(Schindelin et al., 2012; Schneider et al., 2012). Although Imaris (Zeiss) and Neurolucida360 (MBF Bioscience) are proven 3D software tools and generate striking images(Quinta et al., 2015; Wang et al., 2018b), the fact that the code is not-open source is an important issue when developing a method for quantitative analysis. ImageJ requires a lot of random access memory (RAM) power of a computer, but since it is open source and can process data in various programming languages, it is possible to perform not only quantification, but also fiber-tracking and developments of plugins at will.

### 4.4. Future tasks

The scar in the center of the injury remains resistant to current clearing procedures, which complicates the observation of this area of great interest (Supplementary Figure 2). The remaining yellowish coloration of the center of the lesion is thought to originate from the crosslinking of extracellular matrix (ECM) proteins by the hydrogel from the PACT procedure. Pretreatment to digest this ECM prior to the clearing procedure may result in significant improvement, and this should be investigated in a future study.

The ultimate objective of clearing-based analyses of the CNS is to apprehend a detailed three-dimensional understanding of the neural circuits(Bachmann et al., 2014; Chovsepian et al., 2017; Economo et al., 2016; Ye et al., 2016). In the present study, we acquired continuous images of long segments of the spinal cord with high resolution. Nevertheless, a method to track single fibers and comprehensively determine the number and position of their branches or varicosities remains to be developed. By assessing the type of fibers that regenerate within damaged circuits when functional recovery is demonstrated by some treatments, one can understand what allows and what limits functional recovery, and would take an important step toward effective treatments.

BDA was used as a tracer, and staining was performed with fluorescently labeled streptavidin. However, to identify the cell types that interact with the descending circuit, it will be necessary to combine the present method with a immunostaining of block tissue.

## 5. Conclusion

Using a mouse injury model spinal cord, a method for tracing the CST, clearing by the modified Passive Clarity method, observing with a Light-sheet microscope, and performing three-dimensional reconstruction, we have established a methodology that enables straightforward depiction up to single fibers. In the future, it will be necessary to achieve reliable quantitative three-dimensional analyses of the injured spinal cord upon various treatments to ultimately define the most appropriate therapeutic strategy in the different pathological situations.

## Abbreviations

BDA: biotinylated dextran amine
CNS: central nervous system
CST: corticospinal tract
PACT: passive clarity technique
RI: refractive index
SC: spinal cord
SCI: spinal cord injury

## Acknowledgments

We appreciate the assistance provided by S. Shibata, J. Koyama, S. Nori, O. Tsuji, K. Sugai, and S. Kawashima, all of whom are members of the spinal cord research team in the Department of Orthopedic Surgery and Physiology, Keio University School of Medicine, Tokyo, Japan. This work was supported by the Japan Agency for Medical Research and Development (AMED; grant nos. JP13bm0204001 to H.O. and M.N.), and the General Insurance Association of Japan Medical Research Grant to K.N and M.S.

## Declaration of Interest

M.N. declares a consultancy role with K-Pharma Inc. and research funding from RMic and Hisamitsu. H.O. declares a leadership position at Keio University School of Medicine and is a compensated scientific consultant for San Bio Co. Ltd. and K Pharma Inc. These companies have no relationship with the present study. The other authors declare no competing interest.

**Supplementary Figure 1.**
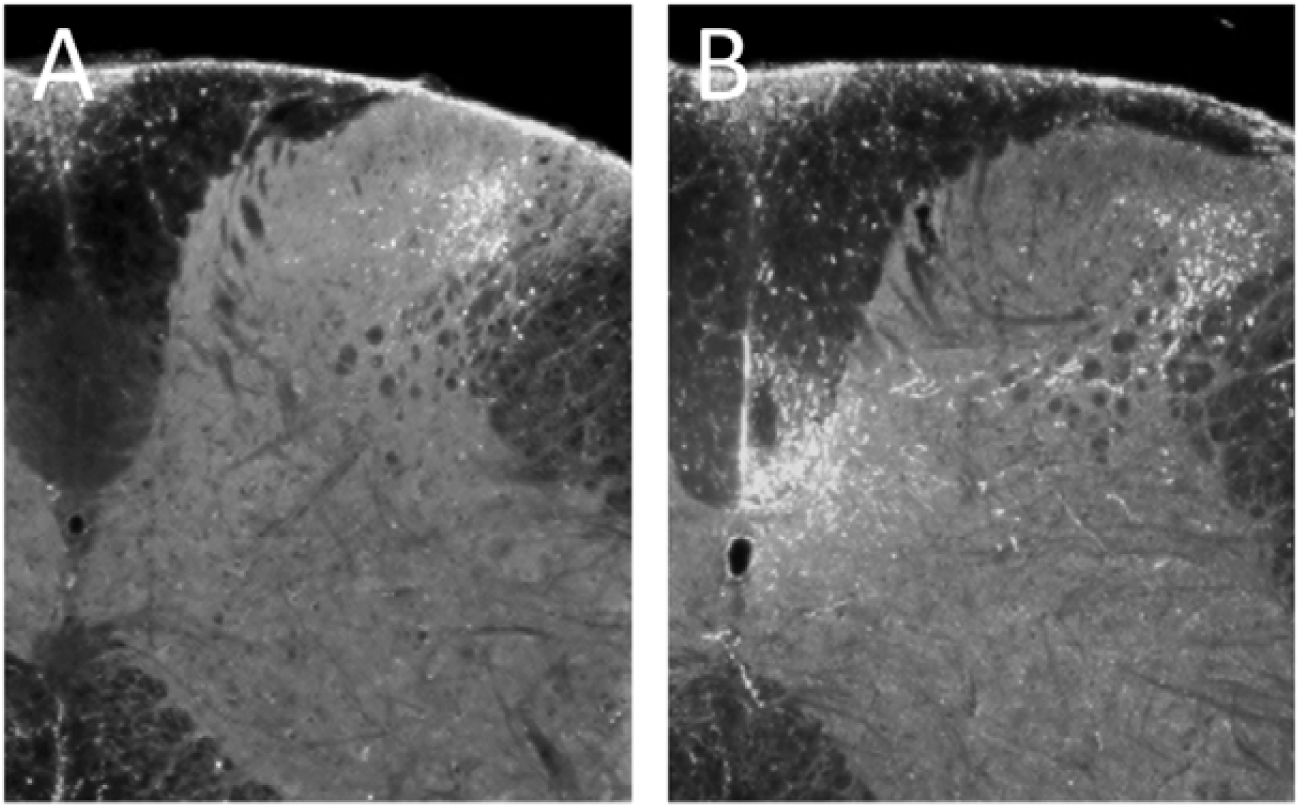
Penetration of the staining solution during the staining process. Comparison of the staining of day 1 (A) and day 7 (B) of the BDA-labeled tissue. On the first day of staining, only the superficial layer was stained, but on the seventh day, the deep white matter was sufficiently stained. Note that the background has not increased even on the seventh day. For this comparison, 4 mm-long spinal cord pieces and axial sections located in the middle (2 mm from the end of the block) were used.

**Supplementary Figure 2.**
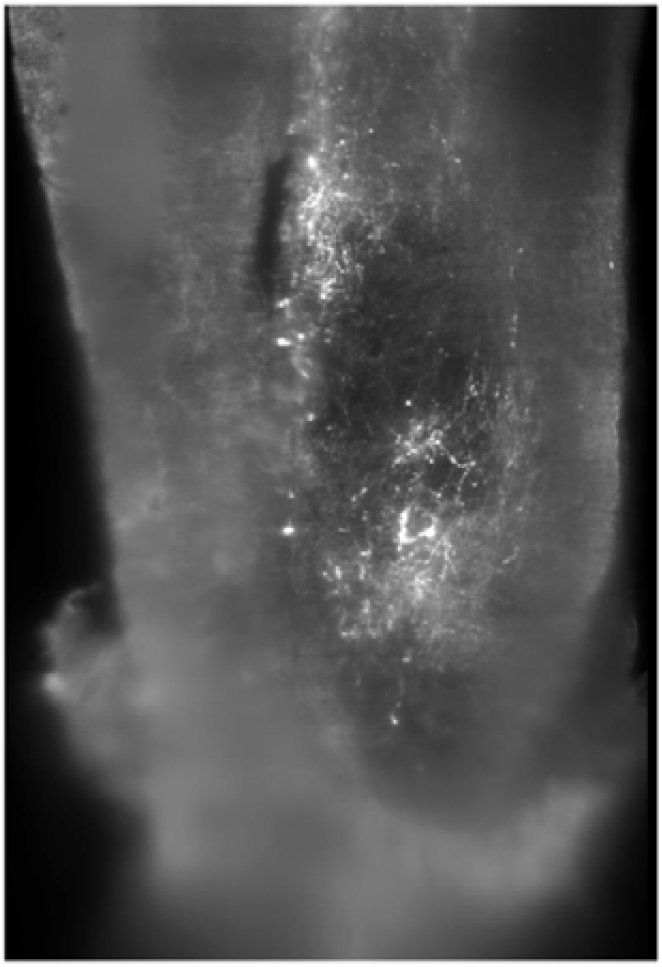
State of the lesion center. Image of the lesion site of the spinal cord. Around the lesion, the resolution of the image decreases, suggesting insufficient clearing of the tissue. This might be related to the remaining extracellular matrix, which negatively affects the permeability of the area for the clearing/staining reagents.

**Supplementary Movies 1-4** Continuous images acquired with a light-sheet microscope. Each movie shows all images captured by light-sheet microscope for each shooting angle. Images were taken continuously from three directions referring to the axes of the spinal cord. See the figure legend in Fig. 4 for imaging conditions. In the axial (1), sagittal (2) and coronal (3) axes, the resolution allows one to clearly delineate individual fibers throughout the entire circumference of the spinal cord. (4) Tiled image in the sagittal direction.

**Supplementary Movies 5-7** Three-dimensional images reconstructed by continuous images. Each movie shows three-dimensional appearance which was reconstructed by ImageJ 3D-project plugin. (5) Reconstruction of sagittal images. (6) Reonstruction of sagittal images with high magnification. (7) Reconstruction of sagittal images tiled in the axial direction of the spinal cord.

